# Brain-inspired Weighted Normalization for CNN Image Classification

**DOI:** 10.1101/2021.05.20.445029

**Authors:** Xu Pan, Luis Gonzalo Sánchez Giraldo, Elif Kartal, Odelia Schwartz

## Abstract

We studied a local normalization paradigm, namely weighted normalization, that better reflects the current understanding of the brain. Specifically, the normalization weight is trainable, and has a more realistic surround pool selection. Weighted normalization outperformed other normalizations in image classification tasks on Cifar10, Imagenet and a customized textured MNIST dataset. The superior performance is more prominent when the CNN is shallow. The good performance of weighted normalization may be related to its statistical effect of gaussianizing the responses.

## 1 Introduction

Normalization layers are widely used in almost all the modern deep neural networks. In convolutional neural networks (CNNs), normalizations fall in two main categories: global and local, according to if the whole spatial dimension is used in normalizing one neuron. Global normalization is motivated by stabilizing the input distribution change (Ioffe & Szegedy, 2015). Local normalization is inspired from neuroscience studies that neurons in visual areas suppress their neighbors (Carandini & Heeger, 2012). Theoretical studies showed that local normalization can benefit the neural computation by reducing statistical redundancies. If linear filters are applied on a natural image, the redundancy of coding is reflected by the super-gaussian marginal distribution. Local normalization models developed in computational neuroscience have been shown to be able to gaussianize the marginal distribution, reduce the statistical dependencies (Schwartz & Simoncelli, 2001; Sinz & Bethge, 2008; Ballé et al., 2016), and better capture neural data (Coen-Cagli et al., 2015; Burg et al., 2020). Local normalizations have achieved promising results in image processing tasks (e.g. denoising and compression) (Ballé et al., 2016; Ballé et al., 2017). However, in CNN based classification tasks, global normalization typically outperforms the local varieties, and there has been only limited work exploring more effective local normalization approaches (Ren et al., 2017; Liao et al., 2016).

In this study, we considered a normalization layer that better captures the understanding of the normalization computation in the brain. It includes two normalization groups: a group of spatially overlapping neurons at the center location, and a group of surround neurons in the same feature map. The normalization weights are set to be trainable, which is updated end-to-end with other parameters in the CNN. We examined the image classification performance in CNNs with different depths against a variety of other global and local normalizations. We showed that weighted normalization outperforms unweighted normalizations and global normalizations in various datasets. We further studied the effect of the various normalizations on the marginal statistics, showing that the weighted normalization results in more gaussianized distributions.

## 2 Related Work

### Global normalization

The batch normalization (BN) is the vanilla version of the global normalization (Ioffe & Szegedy, 2015). It is the default choice of almost all current neural network designs and many benchmark-beating networks still use it (Tan & Le, 2019). The normalization pool of BN is the whole spatial map of a single channel across several inputs, i.e. a mini-batch. Several variations of batch normalization uses batch size of 1, with various design choices of how many channels are used in the normalization pool: instance normalization (IN) (Ulyanov et al., 2016); filter response normalization (Singh & Krishnan, 2020); group normalization (GN) (Wu & He, 2018); and layer normalization (LN) (Ba et al., 2016). The normalization pool of global normalization is unrealistically large for a biological neuron.

### Local normalization

Local normalization only uses local units in the normalization pool, which is closer to the brain computation than global normalizations. Examples of local normalization in CNNs are local response normalization (LRN) (Krizhevsky et al., 2012), divisive normalization (DN) (Ren et al., 2017), local context normalization (Ortiz et al., 2020) and region normalization (Yu et al., 2020). There are at least two properties of computations in the brain that are not captured by these local normalizations, which are apparent in both experimental and theoretical neuroscience studies: 1) the normalization weights, even for neurons with spatially overlapping receptive fields, are not uniform (Burg et al., 2020); and 2) surround normalization is more prevalent for within-channel neuron pairs than neuron pairs in different feature maps (Schwartz & Simoncelli, 2001).

### Weighted normalization

This refers to normalization with weights that are not uniform. From some computational neuroscience studies (Wainwright et al., 2001), to maximize the information content, normalization weights are related to the (co)variance of the normalization pool. Since the spatial dependencies are usually different for different neurons, the normalization weights are rarely uniform. Weighted normalization has been used in tasks such as image denoising, image compression and neural response predictions (Ballé et al., 2016; Ballé et al., 2017; Burg et al., 2020). Ballé et al. (2016) found that a form of weighted normalization can gaussianize the image data. Models with trainable normalization weights can predict neural responses better than uniform competitors (Burg et al., 2020). However, the influence of weighted normalization in image classification with CNNs is poorly understood.

## 3 Methods

In this study, to test the effects of both weighted normalization and incorporating surround normalization, we proposed two normalization models (shown in figure 1). The center-only normalization pool only contains 8 neurons at the same spatial location but from different feature maps (colored orange). Beside the neurons in the center-only pool, the center-surround normalization pool also contains 8 surround neurons in the same feature map as the neuron to be normalized (colored green). Each neuron in the normalization pool has a trainable weight and is constrained to be positive. In other words, the normalization term is the square root of the weighted sum of squared activations:

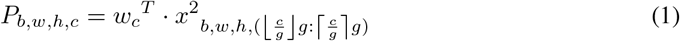

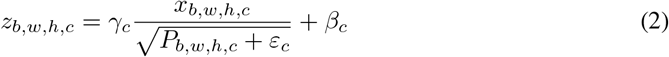

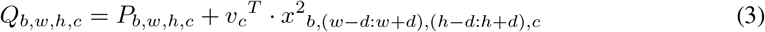

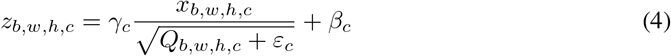

**Figure 1:**
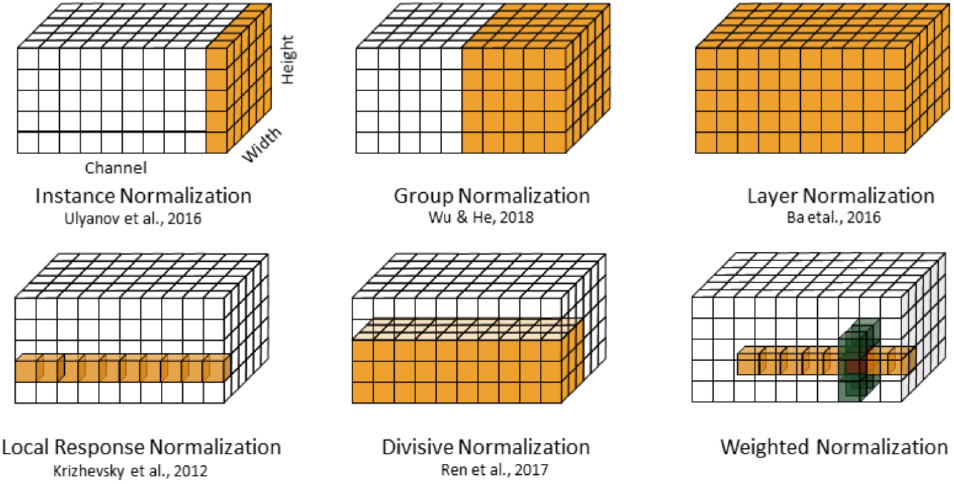
Weighted versus other normalizations. The red neuron is to be normalized. Orange and green units are center and surround pools respectively, with the weights learned during task training.

*P* and *Q* are center and center-surround pool respectively. Subscripts *b,w,h,c* are sample, width, height and channel dimensions. *g* is the group size. We choose group size 8 to match the surround pool. *d* is the surround distance. *w_c_* and *v_c_* are channel-wise normalization weights for center and surround neurons. ε_c_ is a positive channel-wise smoothing factor that can keep the stability of the optimization. Like other normalizations, channel-wise gain *γ* and bias *β* is added after dividing the normalizaiton pool. *z* is the output of the normalization layer. When testing the effect of trainable weights, we compare the trainable model with the one with fixed *w_c_* and *v_c_*. The fixed center-only model differs from LRN by an extra gain γ, bias β and choice of hyperparameters. Both the normalization weighting and the tuned surround are closer to biology than other normalizations used in CNNs. Note that in this study, we are seeking to find when brain-like computations can benefit the task, rather than competing with the benchmark.

## 4 Results

### 4.1 Classification

We hypothesize that our normalization is most beneficial when the network is shallow, and can better benefit from the nonlinearity and its potential for gaussianizing and decoupling the neurons since neurons in deeper layers are inherently more gaussian and less dependent (Ballé et al., 2017; Sánchez Giraldo & Schwartz, 2019). To test this, we first examined image classification performance for the Cifar10 dataset (Krizhevsky & Hinton, 2009) with 6 CNN models with 1 (shallowest) to 6 (deepest) convolution layers. Subsequently, we also considered the AlexNet architecture versus a smaller Alexnet with the Imagenet dataset (Deng et al., 2009), and digit classification for texturized MNIST digits.

The testing accuracy of weighted normalization versus other normalizations for Cifar10 dataset is shown in figure 2. When we reduce the number of layers of the CNN model, weighted normalizations performed better than other normalizations. The effect is most prominent when the CNN has one (or two) convolution layers. Setting weights to be trainable is always better than fixed. Fixing the normalization weights reduced about 2% testing accuracy from the weighted versions. When the CNN model is relatively deep, the weighted normalizations show no improvement over other normalizations, such as BN and IN, but are competitive. Including surround does not further improve, and sometimes reduces the performance, compared to the center-only weighted normalization.

**Figure 2:**
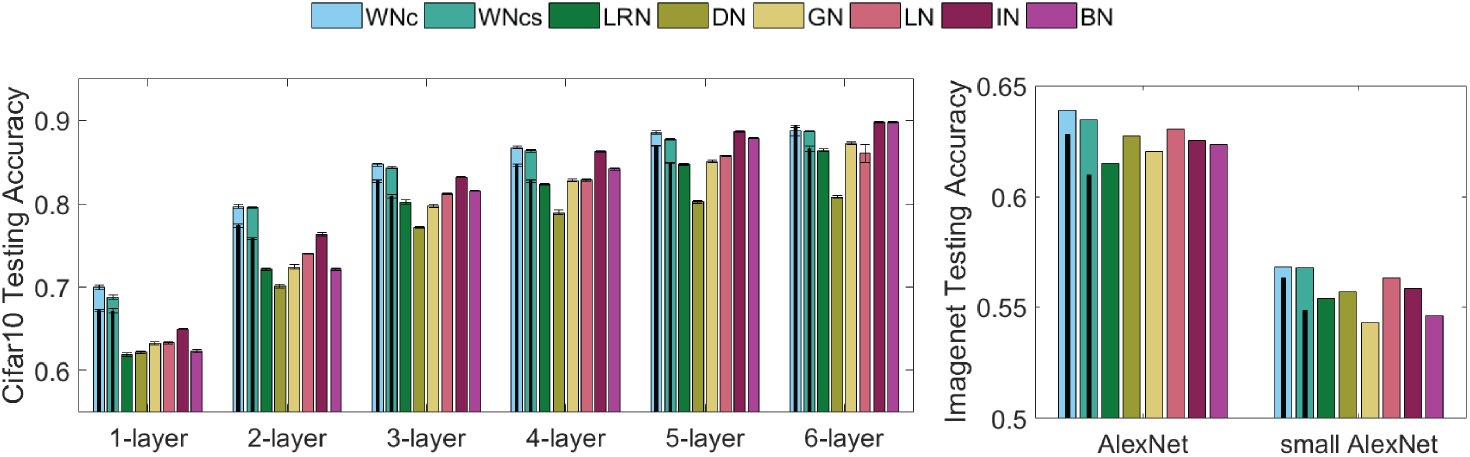
Top-1 testing accuracy for the Cifar10 (left) and Imagenet (right) dataset. WNc is weighted center normalization. WNcs is weighted center-surround normalization. The inner thin black bar is when normalization weights are fixed. Error bar shows standard error of the mean (SEM) from three random initializations

To test if the results of the Cifar10 dataset could generalize to larger models and datasets, we trained Alexnet (5 convolution layers) and “small Alexnet” (3 convolution layers) on the Imagenet dataset. Top-1 testing accuracy is shown in figure 2 right. We observed similar results as in the Cifar10 experiment.

From the Cifar10 and Imagenet experiments, we did not see further improvement when including surround normalization. We hypothesize that this could be because the image classification does not require strong differentiation between center and surround features, so surround normalization failed to benefit and even reduced the fit. A study that showed CNNs are biased to texture information supports our hypothesis (Geirhos et al., 2019). To test this hypothesis, we designed a new dataset (textured MNIST) based on the MNIST dataset, randomly imposing textures from a texture pool to the foreground and background (examples shown in figure 3). In this dataset, texture itself doesn’t provide label information. All the label information are from the outlines of the texture. So for this dataset, weighted center-surround normalization could have better performance, since it has a role of contrasting feature from the context.

**Figure 3:**
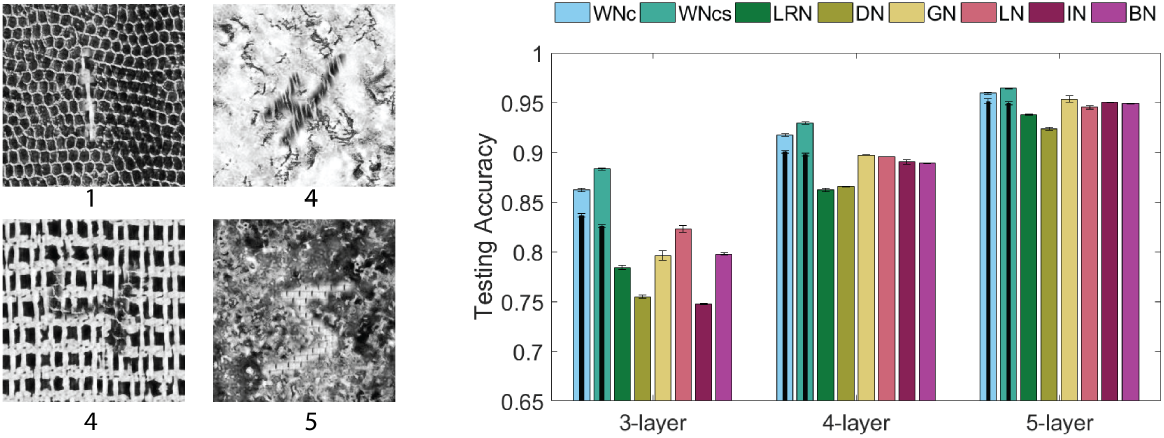
Left: examples of textured MNIST. Right: testing accuracy for textured MNIST dataset.

Testing accuracy is shown in figure 3. Similar to the other experiments, the weighted normalizations outperformed other normalizations. The superiority of weighted normalization is more prominent in shallower CNNs. We further found that incorporating surround normalization led to better performance, particularly for the shallower networks.

### 4.2 Statistics after normalization

We examine our hypothesis that WN induces a more efficient representation by gaussianzing the marginal response distribution. In figure 4, we show that compared to BN which has no effect on the marginal distribution, WNc can better gaussianize the marginal distribution. Already after normalization in the second layer, the marginal distribution is very close to a gaussian distribution. This is reflected in the measure of negentropy (i.e., the Kullback-Leibler divergence between a distribution and the standard normal). The weighted normalization is one of the most effective normalizations to maximize the entropy (figure 4), while some other normalizations, such as LRN, LN and DN, can also do a decent job.

**Figure 4:**
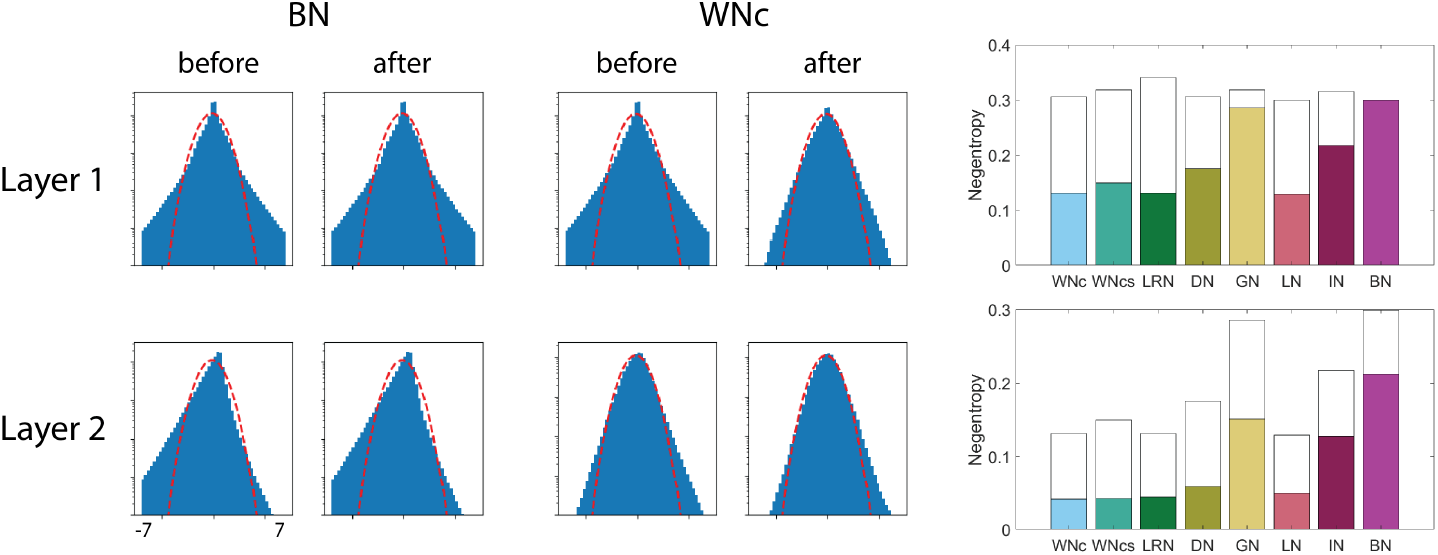
Left: effect of BN and WNc on the mariginal distribution in layer 1 and layer 2 of Alexnet. Red line indicates a standard gaussian distribution. Y axis is in log scale. Right: Negentropy changes at layer 1 and 2. Hollow bars indicate negentropy before normalization.

## 5 Conclusion

We studied a normalization paradigm, weighted normalization, that better reflects the current understanding of the brain. Weighted normalization outperformed other normalizations in image classification tasks with Cifar10, Imagenet and a customized textured MNIST dataset. The superior performance is more prominent when the CNN is shallow. The brain may take advantage of this to form a more efficient representation without having tens of processing stages (i.e. brain areas). We found that the surround pool is important when label information is in the shape of the texture, and expect surrounding information to further be important for other vision tasks. The good performance of weighted normalization may be related to its statistical effect of gaussianizing the responses.

## Acknowledgments

This work was supported by a National Science Foundation Grant 1715475 (to O.S.), and the Scientific and Technological Research Council of Turkey (TUBITAK-BIDEB) 2219 International Postdoctoral Research Scholarship Program (to E.K.).

